# Fadrozole-mediated sex reversal in the embryonic chicken gonad involves a PAX2 positive undifferentiated supporting cell state

**DOI:** 10.1101/2022.08.06.503058

**Authors:** Martin A. Estermann, Craig A. Smith

## Abstract

Gonadal sex differentiation among vertebrates involves divergent fates of a common groups of progenitor cells present in both presumptive ovaries and testes. The first cell type to differentiate gives rise to pre-Sertoli cells in the testis, and pre-follicular cells in the ovary. These cells derive form a common lineage of so-called “supporting cells”. In birds and other egg-laying vertebrates, locally synthesised estrogen has a central role in ovarian development and influence the fate of these supporting cells. Manipulation of estrogen levels during embryonic development induces gonadal sex reversal, providing an experimental setting to evaluate the process of gonadal sex differentiation. Recently, we identified PAX2 as a novel marker of the undifferentiated supporting cell lineage in the chicken embryo, expressed in both sexes prior to overt gonadal sex differentiation. *PAX2* expression is downregulated at the onset of gonadal sex differentiation in both males and females. The analysis of this undifferentiated supporting cell marker, together with Sertoli (male) and pre-granulosa (female) will enhance our understanding of supporting cell differentiation. Here we characterized the supporting cells differentiation process and identified undifferentiated supporting cells in estrogen-mediated sex reversal experiments. Female embryos treated with the aromatase inhibitor fadrozole developed ovotestis, containing pre-granulosa cells, Sertoli cells and PAX2 positive undifferentiated supporting cells. In contrast, male embryos treated with 17β-estradiol showed no PAX2^+^ undifferentiated gonadal supporting cells. Fadrozole time-course as well as multiple dose analysis suggests that supporting cell transdifferentiation involves a dedifferentiation event into a PAX2^+^ undifferentiated supporting cell state, followed by a redifferentiation towards the opposite sex lineage.

## Introduction

Gonadal sex differentiation describes the process of ovary or testis formation among vertebrate embryos. In most species, the gonads differentiate during embryonic or larval life, or shortly thereafter. Among humans, gonadal and external genital differentiation are sometimes atypical. Differences of Sex Development (DSDs) in humans have an incidence of around 1% of all live births. DSDs occurs when the chromosomal, gonadal or anatomical sex are discordant or ambiguous (1, 2). Gonads exhibit a wide range of phenotypes, from total to partial sex reversal, ovotestis or gonadal dysgenesis (3). In some cases, gonads develop normally, and the external genitalia are ambiguous, as in Androgen Insensitivity (AIS) or CAH (Congenital adrenal hypoplasia) (4, 5). Despite the advances genetic diagnosis, many DSD cases lack a definitive molecular diagnosis (6). All current DSDs diagnostical methods do not provide functional information. This is crucial for understanding the etiology and risks associated with specific DSDs mutations, and for clinical management (7). In recent years, animal models have been very instructive in elucidating the molecular genetics of typical and atypical gonadal sex differentiation (8-12).

Comparative analysis of chicken and mouse embryos has demonstrated significant genetic and morphological conservation of gonadal sex differentiation (13, 14). As in human embryos, both chicken and mouse exhibit the same groups of progenitor cell types in the gonad. These are the supporting cells (presumptive Sertoli cells in the testis and granulosa cells in the ovary), steroidogenic cells (presumptive Leydig and theca cells), germ cells (sperm or ova) and other interstitial cells such as vascular progenitors (15-17). This conservation largely extends to the genetic level. *DMRT1, AMH* and *SOX9* are expressed and play decisive roles in testicular morphogenesis, while *WNT4, RSPO1, FOXL2* and *CYP19A1* (aromatase) are important in the ovary (18-21). Therefore, the chicken provides a very tractable model for studying embryonic gonad development (22, 23). In birds and other egg-laying vertebrates, locally synthesised estrogen is required for ovarian development (24-28). The estrogen synthesising enzyme, aromatase, is only expressed in female gonads at the onset of ovarian differentiation (24, 29). Experimental manipulation of estrogen levels during embryonic development has been associated with gonadal sex reversal (30, 31). Inhibition of aromatase enzyme leads to testis or ovotestis formation in genetic females (29, 32-34), while over-expression in genetic males can induce transient ovary formation (35). This model provides an experimental setting to evaluate the process of gonadal sex reversal and gonadal sex diff more broadly.

Recently, we identified PAX2 as a novel marker of the undifferentiated supporting cell population in the embryonic chicken gonad. Furthermore, we found that PAX2 is also expressed in the undifferentiated gonads of all major avian lineages (36). In chicken, PAX2 expression is sharply downregulated at the onset of gonadal sex determination in both males and females (15, 36). The identification of this undifferentiated supporting cell marker, together with Sertoli (male) and pre-granulosa (female) markers now allows for a more complete understanding of the timing and nature of supporting cell differentiation.

Here we characterize the supporting cells differentiation process, and the prevalence of undifferentiated supporting cells in estrogen-mediated sex reversal experiments. Female embryos treated with the aromatase inhibitor, fadrozole, developed ovotestis, containing pre-granulosa cells, Sertoli cells and PAX2 positive undifferentiated supporting cells. In contrast, male embryos treated with 17β-estradiol showed no PAX2^+^ undifferentiated gonadal supporting cells. Fadrozole time-course as well as multiple dose analysis suggests that supporting cell transdifferentiation involves a dedifferentiation phase into a PAX2^+^ undifferentiated supporting cell state, followed by a redifferentiation towards the male (Sertoli) lineage.

## Materials and Methods

### Chicken eggs

Fertilized *Gallus gallus domesticus* HyLine Brown eggs were obtained from Research Poultry Farm (Victoria, Australia) and incubated at 37.5°C under humid conditions. Embryos were staged according to according to Hamburger and Hamilton (37). Experiments were performed in accordance with our institutional animal ethics requirements (Approved Monash University AEC #17643).

### Fadrozole treatment

Eggs were injected at E3.5 (Hamburger and Hamilton stage HH19) (37) with 1mg of fadrozole (Novartis) in 100 µL of PBS or vehicle, as described previously (38). Embryonic urogenital systems were collected at E6.5 (HH30), E9.5 (HH35) or E12.5 (HH38). For double injection experiments, eggs were injected at E3.5 and at E6.5 with 1 mg of fadrozole in 100 µl of PBS or vehicle. Tissues were collected at E9.5.

### 17β-estradiol treatment

Eggs were injected at E3.5 (HH19) with 0.1 mg of 17β-estradiol in 10% ethanol in sesame oil or vehicle, as described previously (38). Tissues were collected at E9.5 (HH35).

### Sexing PCR

Genetic sexing was performed by PCR, as described previously (39). Genetic females (chromosomally ZW) were identified by the presence of a female-specific (W-linked) XhoI repeat sequence in addition to a 18S ribosomal gene internal control. Genetic males (chromosomally ZZ) showed the 18S band only (39).

### qRT-PCR

Quantitative RT-PCR (qRT-PCR) was performed as reported before (36). Briefly, E9.5 (HH35) gonadal pairs were collected and snap frozen in dry ice. After PCR sexing, 3 same-sex gonadal pairs from the same treatment were pooled for each sample, homogenized and RNA extracted using 1mL TRIzol as per the manufacturer’s instructions, (TRIzol, ThermoFisher). Genomic DNA was removed using DNA-free™ DNA Removal Kit (Invitrogen) and 1 μg of total RNA was reversed transcribed into cDNA using Promega Reverse Transcription System (A3500). RT-qPCR was performed using the QuantiNova SYBR® Green PCR Kit. *PAX2* expression levels were quantified by the 2^-ΔΔCt^ method using β-actin as internal control. PCR primers were: *PAX2* Fw: GGCGAGAAGAGGAAACGTGA, *PAX2* Rv: GAAGGTGCTTCCGCAAACTG, *β-actin* Fw: CTCTGACTGACCGCGTTACT and *β-actin* Rv: TACCAACCATCACACCCTGAT. Data was analysed using two-way ANOVA. Significance was determined using Tukey posttest.

### Immunofluorescence

Immunofluorescence of frozen sections was performed as reported previously (15). Briefly, gonadal samples were fixed in 4% paraformaldehyde/PBS for 15 minutes, cryoprotected in 30% sucrose in PBS overnight, embedded in OCT embedding compound and snap frozen at -80°C. 10μm gonadal cryosections were permeabilized with 1% Triton X-100 in 1X PBS for 10 minutes, blocked in 2% BSA in 1X PBS for 1 hour, incubated overnight at 4°C with the primary antibody in 1% BSA in 1X PBS. Primary antibodies used: rabbit anti-PAX2 (Biolegend 901001, 1;500), rabbit anti-DMRT1 (in house antibody; 1:2000), rabbit anti-SOX9 (Millipore AB5535, 1:4000), rabbit anti-Aromatase (in house antibody; 1:4000) and rabbit anti-AMH (Abexa ABX132175; 1:1000). Samples were washed and incubated with the secondary antibody Alexa Fluor 488 donkey anti-rabbit (1:1000) in 1% BSA in 1X PBS. Samples were counterstained with DAPI and mounted in Fluorsave (Millipore). Images were collected on a Zeiss Axio Imager A1 microscope using a Zeiss Axiocam MRc5 camera, using the same exposure time between treatments for expression comparisons. For PAX2 immunostaining, antigen retrieval was performed using the Dako PT Link automated system.

## Results

### PAX2 expression is induced upon fadrozole mediated masculinization

In the chicken embryos, a typical genetic female develops a large ovary, characterised by thickened outer cortex and vacuolated underlying medulla (32, 33). Treatment with a single dose of the aromatase inhibitor Fadrozole results in gonads lacking a cortex and developing testicular cords rather that lacuna in the medulla (33). Here, embryonic day 3.5 (E3.5) chicken eggs were injected with the Fadrozole or vehicle solution (Fig. 1A). Embryos were collected at E9.5 (Fig. 1A), genotypically sexed and *PAX2* gonadal mRNA expression was measured by qRT-PCR. *PAX2* mRNA levels were significantly higher in females treated with fadrozole (FAD), compared with the vehicle control (Fig. 1B).

**Fig 1.**
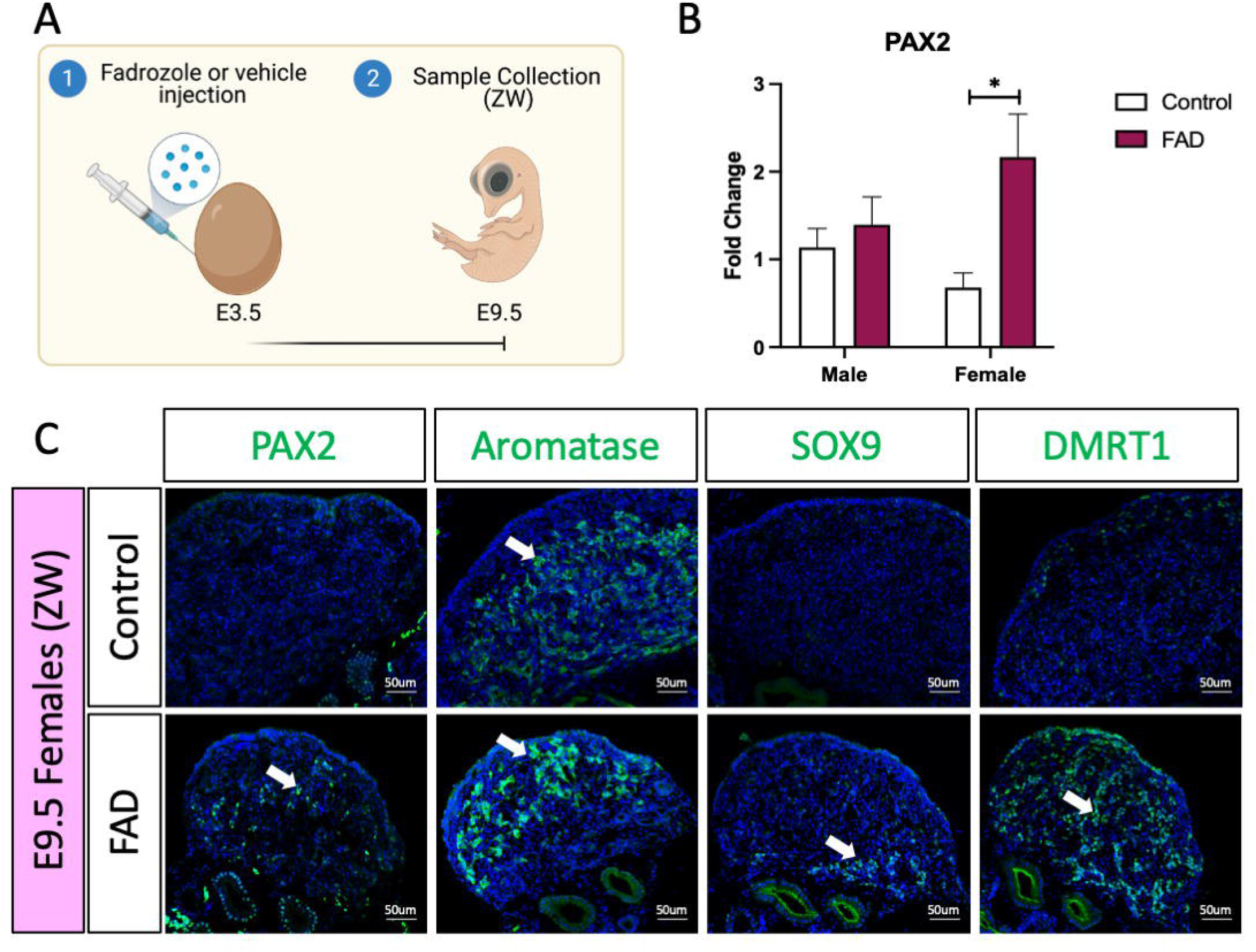
PAX2 expression is induced upon fadrozole mediated masculinization. (A) Schematic figure of the experimental plan. Fadrozole or vehicle solution was injected in chicken eggs at E3.5. Samples were collected at E9.5. (B) *PAX2* qRT-PCR was performed in E9.5 gonadal samples of fadrozole (FAD) or vehicle (Control) treated embryos. Expression level is relative to β-actin and normalized to the male control. Bars represent mean±s.e.m., n=6. * adjusted P<0.05. 2-way ANOVA and Tukey’s post-test. (C) Immunofluorescence against PAX2, aromatase, SOX9 and DMRT1 in E9.5 female (ZW) gonads treated with fadrozole (FAD) or vehicle solution (Control). White arrows indicate positive cells.

In males, *PAX2* expression levels remained unchanged upon treatment (Fig. 1B). To validate the qRT-PCR results, immunofluorescence against PAX2, aromatase (female marker) and male markers SOX9 and DMRT1 was performed in E9.5 female (ZW) gonads treated with fadrozole (FAD) or vehicle (Control). As expected, control left ovaries were larger, expressing aromatase in the medulla but no PAX2, SOX9 or DMRT1 (Fig. 1C). In contrast, fadrozole-treated female left gonads lacked a cortical compartment, and both male (SOX9) and female (aromatase) positive supporting cells co-existed in the same gonad, but in separated defined regions (Fig. 1C). Aromatase positive cells were located in the apical region of the gonad whereas SOX9 positive cells were detected more basally, adjacent to the to the mesonephric kidney. Testis-associate DMRT1 was up-regulated throughout the medulla. PAX2^+^ cells were identified in the gonadal medulla, between SOX9 and aromatase positive supporting cells (Fig. 1C). Taking all together, these results indicate that fadrozole mediated masculinization results in an increase in gonadal PAX2^+^ expression among cells of the female gonad. Based on our previous data, showing PAX2 to be a marker of prior to differentiation, these cells are interpreted to be undifferentiated supporting cells.

### PAX2 *expression is not induced upon estrogen-induced feminization*

To evaluate if PAX2 positive undifferentiated cells were also present in estradiol-mediated male to female sex reversed gonads, E3.5 chicken eggs were injected with 17β-estradiol (E2) or vehicle solution (Fig. 2A). Embryos were collected at E9.5 (Fig. 2A), genotypically sexed and *PAX2* expression was measured by qRT-PCR. No significant differences were found in *PAX2* mRNA expression levels, between treated (E2) and control gonads in both sexes (Fig. 2B). PAX2 Immunofluorescence was performed on E9.5 male (ZZ) gonads treated with 17β-estradiol or vehicle, but no PAX2 positive cells were detected in the gonadal medulla (Fig. 2C), consistent with the qRT-PCR results. To evaluate if, in fact, the gonads were sex reversed, aromatase, SOX9 and DMRT1 protein expression were evaluated by immunofluorescence. As expected, control male gonads showed no aromatase expression and medullary expression of SOX9 and DMRT1 (Fig. 2C). In contrast, E2 treated gonads showed higher ectopic activation of aromatase and lower levels of the male markers, SOX9 and DMRT1 in the medulla (Fig. 2C). Additionally, DMRT1 positive germ cells were detected in the cortical region of the gonad, as is seen in typical left ovaries. These results indicate that 17β-estradiol mediated sex reversal did not induce PAX2 positive undifferentiated supporting cells at E9.5.

**Fig 2.**
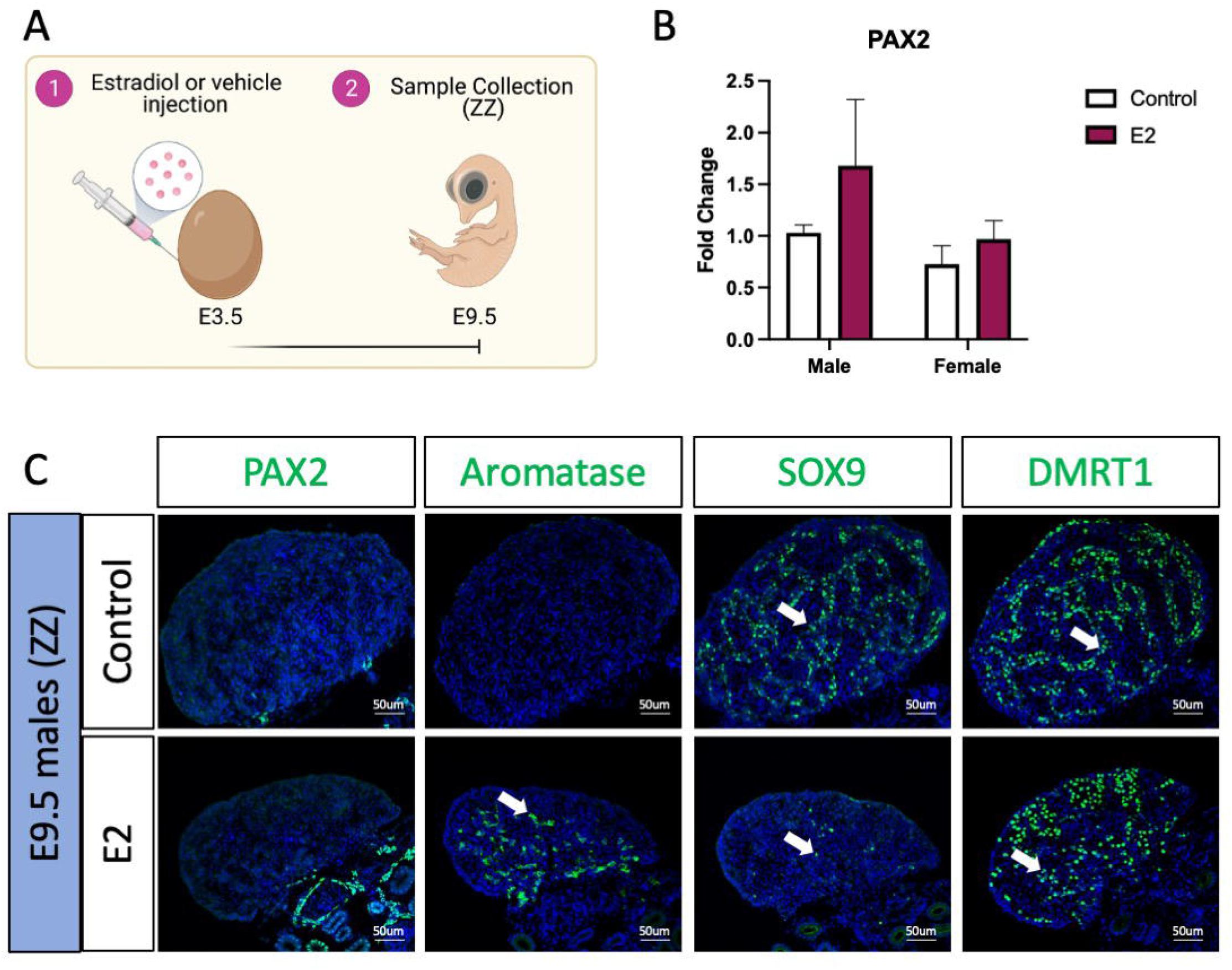
PAX2 expression is not induced upon estrogen-induced feminization. (A) Schematic figure of the experimental plan. 17β-estradiol or vehicle solution was injected in chicken eggs at E3.5. Samples were collected at E9.5. (B) *PAX2* qRT-PCR was performed in E9.5 gonadal samples of 17β-estradiol (E2) or vehicle (Control) treated embryos. Expression level is relative to β-actin and normalized to the male control. Bars represent mean±s.e.m., n=6. 2-way ANOVA and Tukey’s post-test. (C) Immunofluorescence against PAX2, aromatase, SOX9 and DMRT1 in E9.5 male (ZZ) gonads treated with 17β-estradiol (E2) or vehicle solution (Control). White arrows indicate positive cells.

### PAX2 *expression is not altered in E6*.*5 and E12*.*5 in fadrozole mediated masculinization*

To evaluate if fadrozole treatment in ZW embryos resulted in an increase of undifferentiated supporting cells throughout gonadal development, fadrozole was injected in E3.5 eggs and gonads were collected at E6.5 and E12.5 (Fig. 3A). Samples were genetically sexed and ZW gonads were immunostained for male (SOX9 and DMRT1), female (aromatase) and undifferentiated (PAX2) supporting cell markers. At E6.5, PAX2 was detected in both control and FAD-treated embryos. Some numbers of Aromatase^+^ cells were also detected, indicating the onset of ovarian differentiation. However, the testis determining factor DMRT1, showed clear upregulation in the female gonad treated with fadrozole aromatase inhibitor. By E12.5, PAX2 expression had been extinguished suggesting that by this developmental stage, all the supporting cells committed to a pre-granulosa or Sertoli cell fate (Fig. 3C). Control gonads showed a typical ovarian structure, with aromatase^+^ vacuolated medulla and an enlarged cortex, containing DMRT1 positive germ cells, and an aromatase positive medulla (Fig. 3C). As expected, E12.5 fadrozole treated gonads lacked a properly defined cortex (Fig. 3C). Aromatase positive pre-granulosa cells located extensively throughout the gonad, whereas SOX9^+^ DMRT1^+^, presumably Sertoli cells, are in the basal region of the gonad (Fig. 3C (Fig. 1C). This data suggests that by E12.5, fadrozole treated gonads at E3.5 reverted to an ovarian phenotype.

**Fig 3.**
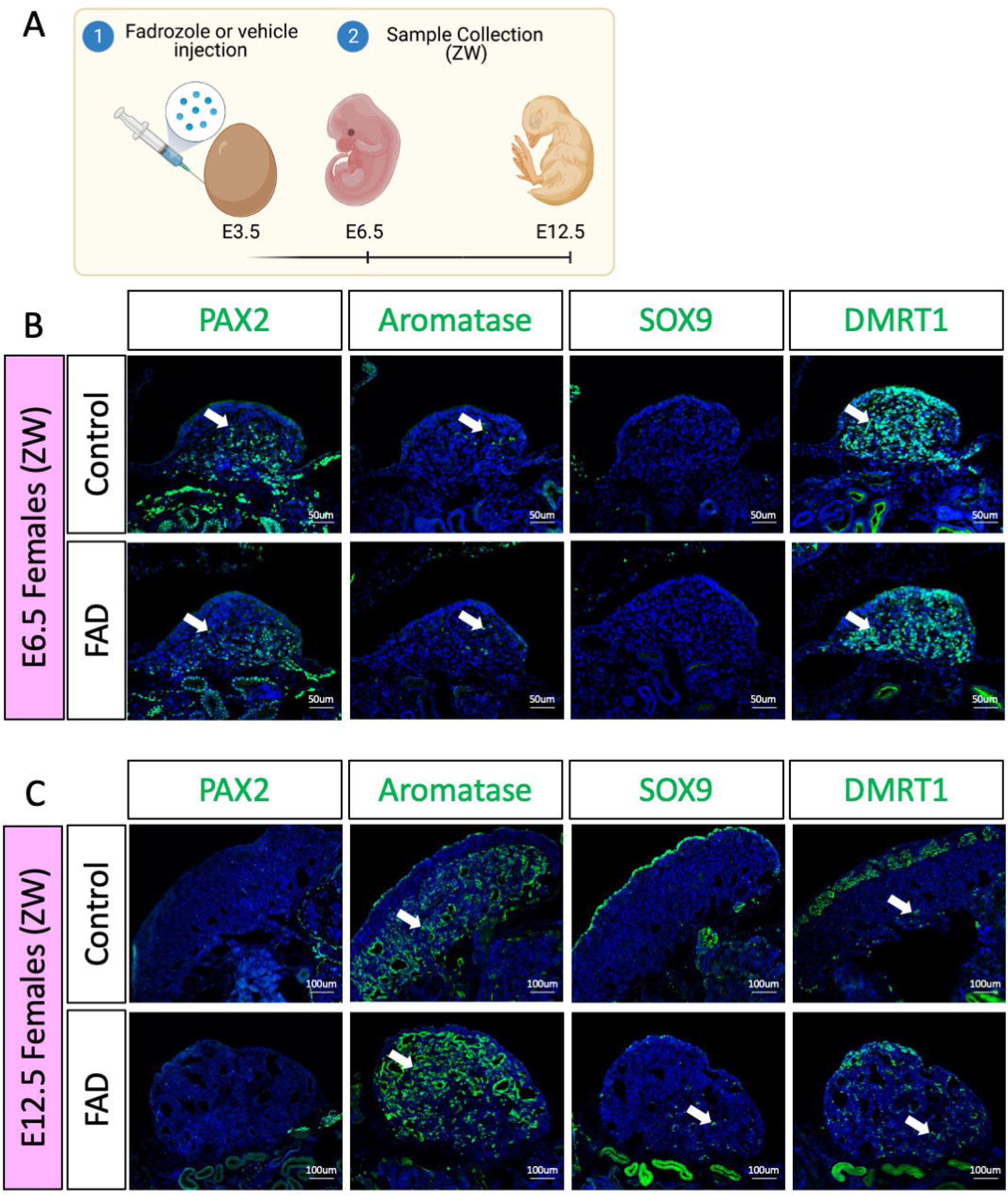
PAX2 expression is not altered in E6.5 and E12.5 in fadrozole mediated masculinization. (A) Schematic figure of the experimental plan. Fadrozole or vehicle solution was injected in chicken eggs at E3.5. Samples were collected at E6.5 or E12.5. (B) Immunofluorescence against PAX2, aromatase, SOX9 and DMRT1 in E6.5 female (ZW) gonads treated with fadrozole (FAD) or vehicle solution (Control). (C) Immunofluorescence against PAX2, aromatase, SOX9 and DMRT1 in E12.5 female (ZW) gonads treated with fadrozole (FAD) or vehicle solution (Control). White arrows indicate positive cells.

### PAX2 *undifferentiated supporting cells are lost upon booster injection of fadrozole*

As fadrozole treatment at E3.5 only results in changes in PAX2 expression pattern at E9.5, but not in E6.5 or E12.5, we wanted to know if those PAX2 undifferentiated cells were a consequence of fadrozole decay over time. To address this question, a double dose experiment was performed, where eggs were injected at E3.5 and E6.5 with vehicle solution (control) or fadrozole, generating 4 different conditions (Fig. 4A). Embryos were collected at E9.5 and immunofluorescence for AMH, SOX9, DMRT1, PAX2 and aromatase was performed (Fig. 4B). As expected, control ovaries expressed aromatase but no PAX2, SOX9, DMRT1 or AMH. Similar to the results from figure 1C, a single dose of fadrozole at E3.5 resulted in gonads expressing male supporting markers (DMRT1, SOX9, AMH), female markers (aromatase) and undifferentiated PAX2 positive cells (Fig. 4B). In contrast, single dose of fadrozole at E6.5 resulted in no PAX2 expression in the gonads, but both Sertoli and pre-granulosa cells coexisting in the gonadal medulla (Fig. 4B). These results indicate that PAX2 upregulation is not a direct result of fadrozole treatment, and that the timing of fad administration is important. Moreover, a similar phenotype was seen in gonads treated with double doses of fadrozole at E3.5 and E6.5 (Fig. 4B). Despite both male and female markers coexisted in the gonadal mesenchyme, no PAX2 expression was detected (Fig. 4B). This suggests that despite one dose at E3.5 results in PAX2 expression, a booster dose at E6.5 inhibits PAX2 expression at E9.5. Taking all together, these results suggests that the undifferentiated supporting cells present at E9.5 are not a direct result of fadrozole injection. Instead, this PAX2 undifferentiated population could be a result of the decay or metabolism of fadrozole at E9.5. The lack of PAX2 positive cells in the gonadal medulla when a booster dose of fadrozole is injected E6.5 supports with this idea. The lack of functional fadrozole at E9.5 results in disinhibition of aromatase and the production of estrogen. Estrogen then could induce the differentiation into pre-granulosa cells while inhibiting Sertoli differentiation. The upregulation of PAX2 suggests that Sertoli to pre-granulosa trans-differentiation involves a dedifferentiation into undifferentiated PAX2^+^ supporting cells, followed by a redifferentiation towards pre-granulosa cells.

**Fig 4.**
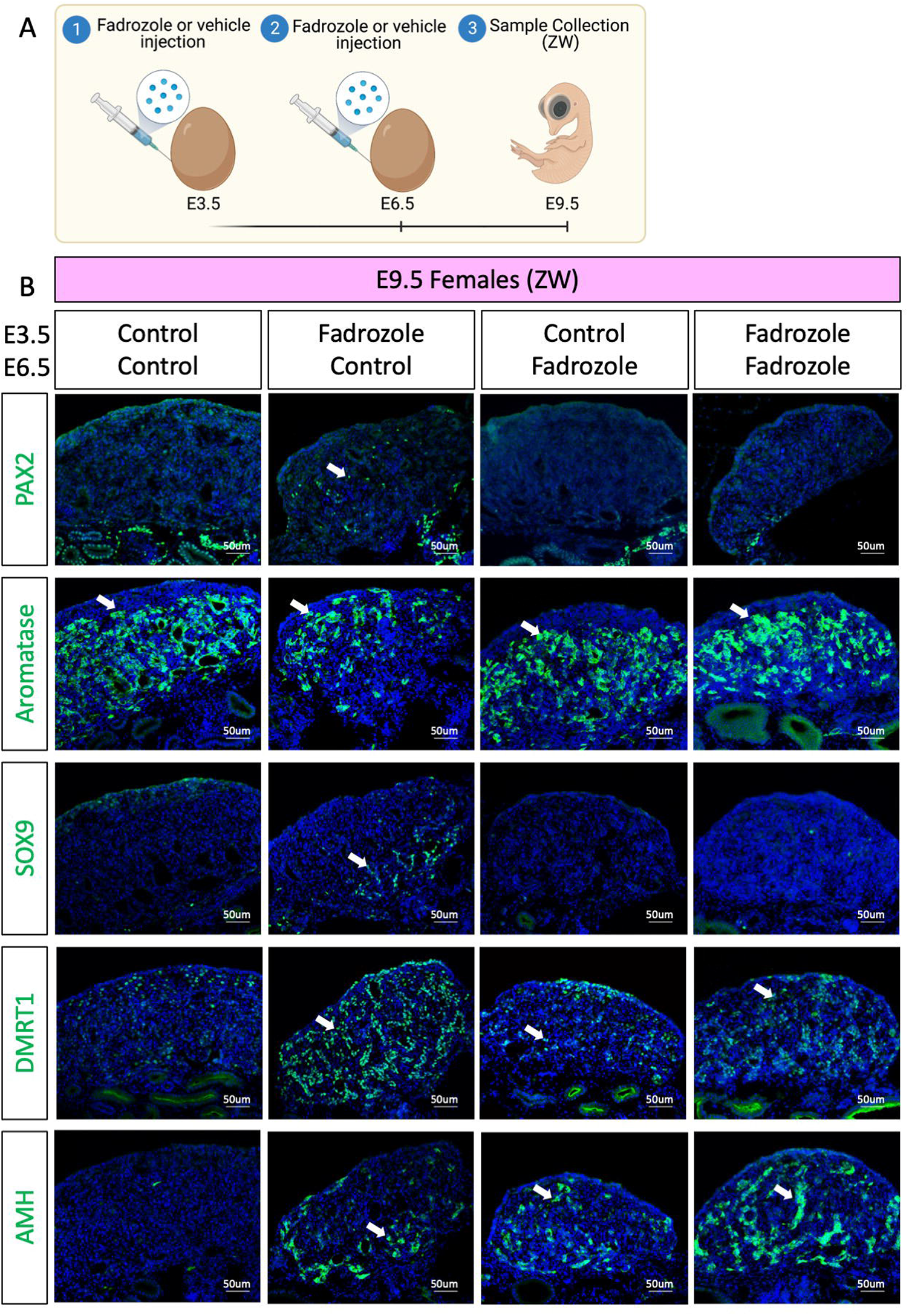
PAX2 undifferentiated supporting cells are lost upon booster injection of fadrozole. (A) Schematic figure of the experimental plan. Fadrozole or vehicle solution was injected in chicken eggs at E3.5 and E6.5. Samples were collected at E9.5. (B) Immunofluorescence against PAX2, aromatase, SOX9 and DMRT1 and AMH in E9.5 female (ZW) gonads treated with fadrozole or vehicle solution (Control). White arrows indicate positive cells.

## Discussion

Gonadal sex reversal occurs when there is a discordance between the genetic/chromosomic and gonadal sex (40). Despite the genetic sex is determined at fertilization, the gonads are sexually determined later during embryonic development (6, 41). Initially, both male and female gonads develop similarly (42). These undifferentiated gonads have the potential of becoming a testis or an ovary, depending on the genetic or environmental signals they receive (43). Among the genetic signals, sex chromosome linked genes like *SRY* in humans or *DMRT1* in birds are sufficient and necessary for testicular development (44-46). Misexpression of those genes in chromosomal females results in testicular development (19, 47). Additionally, *SRY* translocation from the Y to the X chromosome during male meiosis is the cause of the majority of the 46,XX DSD cases (48).

Among the estrogen plays a strong role in ovarian differentiation. Modulation of estrogen levels resulted in gonadal feminization in birds, reptiles, and eutherian mammals (25, 29, 31, 49-51). In placental mammals such as mice, estrogen seems to have a role in gonadal maintenance, as estrogen receptor α and β double knock out result in postnatal gonadal sex reversal (52). In birds, exogenous modulation of estrogen levels can influence the embryonic gonadal fate, independent of the genetic sex. Injection of fadrozole, an estrogen synthesis inhibitor, in female embryos results in testicular development, upregulating *DMRT1* and *AMH* and downregulating *FOXL2* and *aromatase* (29, 34, 38, 53). In some cases, these gonads reverted to ovotestis, upregulating aromatase at around day 7 to 9 of development (32, 38, 54). Our results agree with these reports, showing both aromatase and SOX9 expression in the same gonads at E9.5 and E12.5, upon fadrozole treatment (Fig. 1C and 3C). Surprisingly, aromatase-expressing embryonic pre-granulosa cells were confined to the apical region of the gonad whereas the SOX9 positive Sertoli cells were present at the base of the gonad, suggesting that some cell populations are more susceptible to redifferentiation. Two main Sertoli cell types were identified in chicken embryonic testis (15). One population located in the most apical region of the gonad expressing low levels of Sertoli markers *SOX9* and *DMRT1*, and the other located more basally, expressing higher levels of *SOX9* and *DMRT1* (15). *DMRT1* is required to maintain the Sertoli cell identity in testis (46, 55, 56). Lower *DMRT1* expression in the apical region could explain the higher susceptibility of these supporting cells to transdifferentiate. On the other hand, a higher sensitivity to estrogens could be also responsible to this phenomenon, although this remains unexplored. Further research is required to evaluate why the apical supporting cell population is more sensitive to transdifferentiation than the basal one.

The process of supporting cell re-differentiation coincides with the upregulation of PAX2 in the gonads, a marker only expressed in undifferentiated supporting cells (15, 36). This suggests that Sertoli cells should revert into a more undifferentiated or bipotential state before they re-differentiate into pre-granulosa cells. These PAX2^+^ “undifferentiated” supporting cells are located in between the SOX9^+^ Sertoli and the aromatase positive pre-granulosa cells. In transdifferentiating adult murine gonads, cells expressing both FOXL2 and SOX9 are detected, suggesting the presence of intermediate double positive cell states (55). Further research is required to confirm that if in fact these are three independent populations or if double or triple positive cells are detected. PAX2, SOX9 and aromatase or FOXL2 immunolabelling in the same gonadal sections will be crucial to address these inquiries.

It is curious that PAX2 was upregulated only during fadrozole mediated but not in estrogen mediated sex reversal. This could suggest that female to male sex reversal occurs differently than male to female sex reversal, perhaps due to the early elevated expression of DMRT1 is males that prevents such de-differentiation. Another possibility is that we missed the PAX2 expression / undifferentiated window in our time series. In fadrozole mediated sex reversal, PAX2 was differentially expressed at E9.5 but not at E6.5 or E12.5. This suggest that this process occurs only at a short period of time. Only E9.5 gonads were evaluated in 17β-estradiol mediated sex reversal. Further research should expand our analysis to other embryonic stages before and after E9.5 to elucidate if pre-granulosa to Sertoli re-differentiation also involves an undifferentiated PAX2^+^ state. Similarly, intermediate states between E6.5 and E12.5 in fadrozole mediated sex reversal are required to precisely define the timeframe of this dedifferentiation and redifferentiation process. This would expand our knowledge in this poorly studied mechanism.

The chicken model is an ideal system to study genetic, environmental, and hormonal factors that control normal or abnormal gonadal differentiation. Estrogenic mediated sex reversal provides a valuable tool to model and study DSDs, especially partial sex reversion and ovotestis development.

## Acknowledgments

The authors acknowledge use of the facilities and technical assistance of Monash Histology Platform, Department of Anatomy and Developmental Biology, Monash University.

## Data Availability

All data generated or analysed during this study are included in this published article.

